# Hybrid Neural Network Models Explain Cortical Neuronal Activity During Volitional Movement

**DOI:** 10.1101/2025.02.20.636945

**Authors:** Hongwei Mao, Brady A. Hasse, Andrew B. Schwartz

## Abstract

Massive interconnectivity in large-scale neural networks is the key feature underlying their powerful and complex functionality. We have developed hybrid neural network (HNN) models that allow us to find statistical structure in this connectivity. Describing this structure is critical for understanding biological and artificial neural networks.

The HNNs are composed of artificial neurons, a subset of which are trained to reproduce the responses of individual neurons recorded experimentally. The experimentally observed firing rates came from populations of neurons recorded in the motor cortices of monkeys performing a reaching task. After training, these networks (recurrent and spiking) underwent the same state transitions as those observed in the empirical data, a result that helps resolve a long-standing question of prescribed vs ongoing control of volitional movement. Because all aspects of the models are exposed, we were able to analyze the dynamic statistics of the connections between neurons. Our results show that the dynamics of extrinsic input to the network changed this connectivity to cause the state transitions. Two processes at the synaptic level were recognized: one in which many different neurons contributed to a buildup of membrane potential and another in which more specific neurons triggered an action potential. HNNs facilitate modeling of realistic neuron-neuron connectivity and provide foundational descriptions of large-scale network functionality.

## 1 Introduction

Artificial neural networks (ANN) are now used to describe results of experiments in which neural activity is recorded during volitional tasks [1–5]. Typically, an ANN is used to make an indirect comparison between global activity patterns common to the network and the empirical data. Here, we take a different approach by developing a hybrid neural network (HNN) composed of neurons with responses matching those recorded from behavioral experiments embedded in a layer of intrinsic artificial neurons. Since the HNN is trained to replicate actual neural activity, it is constrained to have physiological plausibility. Once trained, these networks can be examined to reveal the connectivity structure underlying the activity of the neurons in the behavioral sample. We applied this approach to model neural activity recorded during volitional movement. Monkeys made reaches in different directions as activity from motor cortical neurons was recorded with microelectrode arrays. The recorded populations of single-unit activity were characterized as patterns of neuron-neuron correlation, which were then used to establish realistic extrinsic neural input to the HNNs. Analyses of the recorded data and of the population dynamics show that the cortical activity during single reaches was undergoing rapid changes in state, defined here as a network mode corresponding to a pattern of correlation. Our findings refute the idea that volitional movements are completely predetermined and show how extrinsic input can drive the network into different states. Because the details of the network are fully exposed in these models, we were able to show how structured temporal integration of synaptic potentials changes during behavior to drive state transitions.

## 2 Results

### 2.1 Empirical Data

We recorded simultaneous single-unit activity using Utah arrays in two monkeys as they performed a center-out reach task while viewing a virtual reality monitor (Methods). Simultaneous recordings were made from 67 units in the primary motor cortex of Monkey C and 78 units in the dorsal premotor cortex of Monkey N. Three-dimensional hand position data from Monkey C were recorded at 60 Hz, and those from Monkey N at 100 Hz. Together, these form the database for this study (Methods).

#### 2.1.1 Discrete epochs in firing rates of individual neurons during reach

Directional tuning during reaching has been shown to be non-stationary [6–11], as tuning functions can change in a subset of neurons during a reach, especially as the movement begins. Recent work [12, 13] shows that while these tuning functions may change episodically during the reach, they are stable within each episode. Motor cortical neurons usually have one or two modulation episodes during a reach [12], although we and others [7, 8] have found examples of neurons with three. We use Unit 59 recorded from Monkey C (also shown in [12]) as a three-epoch template (Fig.1A). This neuron, like others with multiple modulation epochs during a reach, has a tuning function that changed rapidly between epochs (Fig.1B). The amplitude of the modulation profile in each epoch is cosine-tuned (Fig.1C). These modulations are well fit with Gaussian profiles, and across neurons, their peaks are confined to one of three epochs during the reach [12]. This temporal consistency suggests that these modulations are common to the population *en masse*. The population-wide consistency of neuronal modulation and the associated changes in tuning suggest that epoch-specific signaling acts as a common factor across neurons responsible for these changes. This can be substantiated using analytical methods to extract separate patterns of correlation in the population associated with these factors.

**Figure 1.**
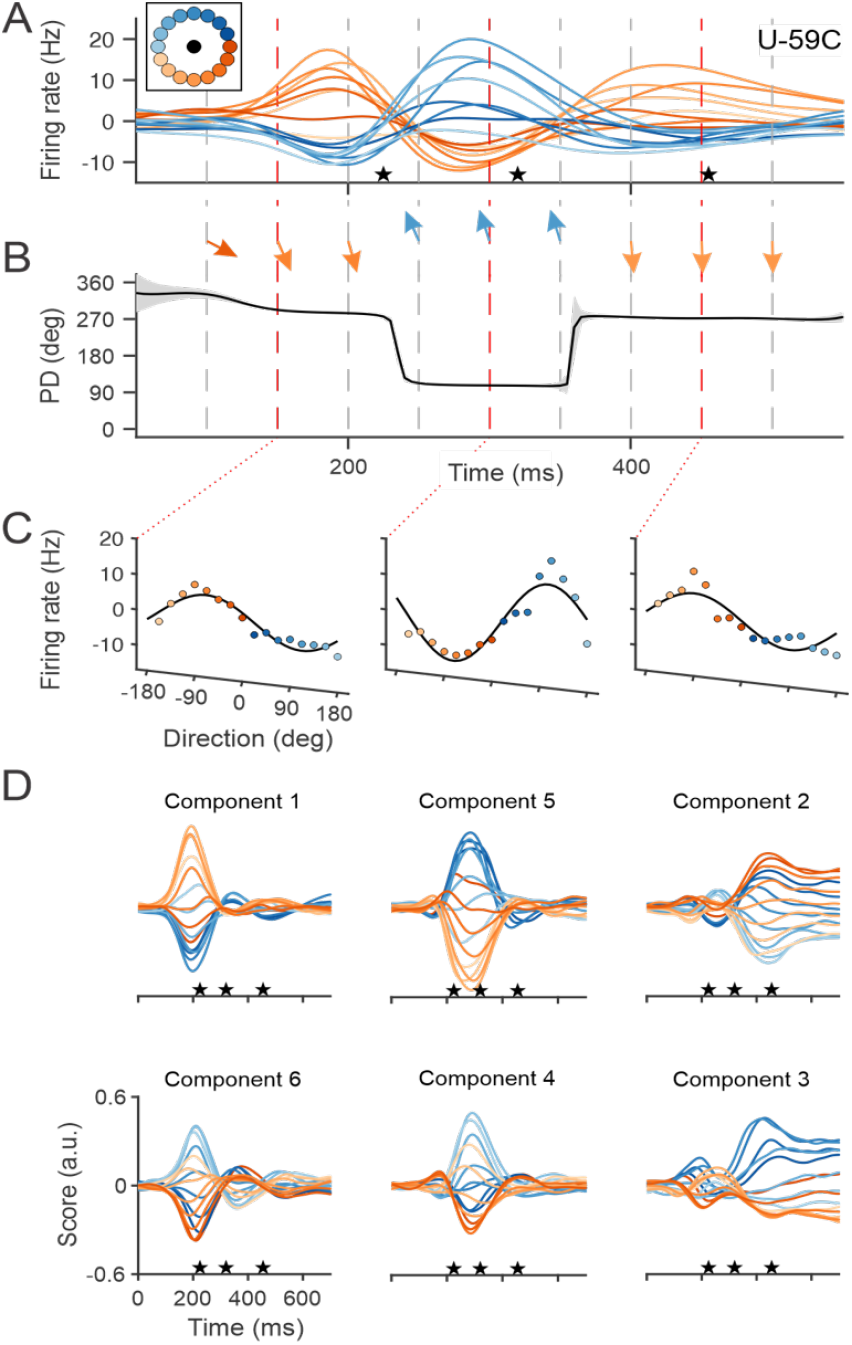
Multiple states in a single reach. **(A)** Trial-averaged firing rates from a single motor cortical unit, U-59C (Unit 59, Monkey C) during reaching in 16 different directions (color-coded inset). The mean firing rate across targets at each time point (non-directional component) was removed. **(B)** Preferred directions changed episodically. Preferred directions are also symbolized by the arrows. **(C)** Tuning functions at different time points (150, 300, 450 ms) during the reach. Firing rates are cosine-fit to the data. **(D)** rPCA scores from Monkey C. The six highest scores are plotted after Promax rotation. Scores are color-coded by movement direction. The profiles in each column have peaks at about the same time in the trial. The stars show the average time of movement onset, peak velocity, and movement completion, respectively.

#### 2.1.2 Correlational structure between neurons reveals successive states

Principal components analysis (PCA) is commonly used for describing patterns of correlation between the activity of neurons in networks [14]. Standard PCA optimizes components that maximize the variance accounted for across the entire data set, which can reduce the explanatory value of the analysis. We applied a rotation to this standard analysis (rPCA – Methods, Fig. S3) which captures individual patterns of correlation. The projection of the data points onto each axis is the score of the component and can evolve through time (Fig. S3B, C).

Consistent with the framework used in network science [15–20], correlation patterns found with rPCA can be considered to represent different states within a population. We found that six PCA components account for 80% of the variance in population firing rate for Monkey C (86% for Monkey N). The rPCA scores from Monkey C are shown in Fig.1D. The single-peak amplitude of each component across movement directions is cosine-tuned. Pairs of scores peaking at the same time correspond to the same network state and are plotted in the columns. The timing and shape of the profiles tend to match those of the modulated firing rates of single neurons [12] and align with movement onset, peak velocity, and target acquisition. The same results were found for data collected from Monkey N (Fig. S4).

### 2.2 Hybrid Network models

We built two HNNs, one with recurrent connectivity and the other with simple feedforward architecture. The latter model was a spiking neural network. Both were trained to reproduce the firing rates of the individual cortical neurons recorded in our center-out experiments. We assumed that the cortical population we sampled was driven by common input with Gaussian profiles resembling those of the rPCA scores (e.g., Fig.1D). We divided the input neurons into three groups, one for each episode of modulation, with their peaks aligned to the start of movement, peak speed, and target acquisition in each trial. Each group consisted of directionally tuned neurons with preferred directions evenly distributed around a circle. This type of input is plausible since many projections to the motor cortex are cosine-tuned to movement direction [21].

This input to the network is *extrinsic* in contrast to the *intrinsic* neuron-neuron connectivity within the network. Both are included in the classic equation for a dynamical system (Eq.1):

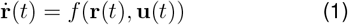

Where:

**r** is a vector composed of neuronal firing rates;

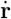 is the change in firing rates over time;

**u** is the external input;

*f* describes how intrinsic and external input contributes to the change of firing rate.

We were able to separate the effect of intrinsic and extrinsic input on our observed neural activity using a recurrent HNN (Fig.2) and found that the extrinsic input was primarily responsible for imparting the modulated firing in our data. To further substantiate this finding, we applied the modeled extrinsic input to a spiking HNN with only feedforward connections to neurons trained to have the same firing rates as those in our data. This network, with only input and output neurons, also replicated our data. Analysis of this network showed that a combination of specific input neurons contributed to each state. Changes in network state were driven by extrinsic input.

**Figure 2.**
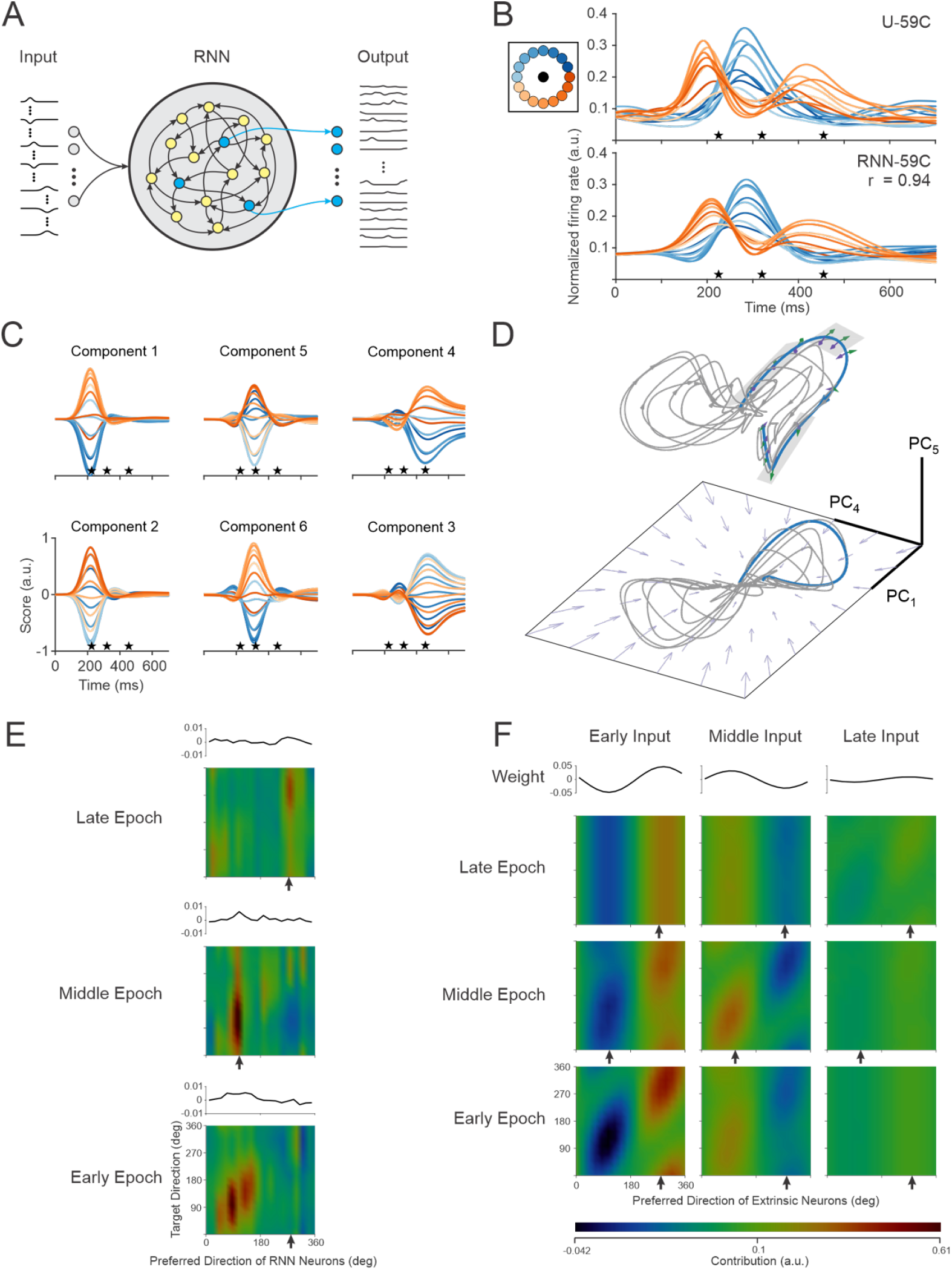
Recurrent neural network. **(A)** Network schematic-Input consists of three groups of 18 directionally tuned neurons with Gaussian temporal profiles corresponding to the rPCA scores in Fig. 1D. Neurons (nodes in blue) in the recurrent layer are trained as output units to generate the empirical firing rates of the recorded neurons in the data set. **(B)** Trial-averaged firing rates of Unit 59, Monkey C. Actual (top, U-59C) and modeled (bottom, RNN-59C) firing rates. Traces begin at target onset. Stars correspond to movement onset, peak speed, and the end of movement. **(C)** Six highest rPCA scores of RNN units after training. **(D)** Neural dynamics of RNN in a 3-D subspace (RNN State Space Trajectory-Suppl. Material). A neural trajectory is plotted for each of the 16 reach directions trials (target onset to target hold). Trajectory components are illustrated with purple (recurrent activation) and green (activation from extrinsic neurons) arrows. Arrows on the plane indicate the change in network activity driven by intrinsic activation at each point on the manifold. **(E, F)** Epoch- and movement-specific contributions to the firing of Unit RNN-59C. For each epoch, the RNN units are divided into 18 groups, according to their preferred direction (PD) in that epoch. The mean weights of the input group are plotted above each heat map. Each pixel of the heatmap shows the total contribution from the units in a PD input group during an epoch of all trials with the same target location. The arrows below each panel designate the preferred direction of the output unit in the respective epoch. **(E)** Intrinsic contributions from the recurrent units to the modeled unit. Contributions of similar magnitude tend to form vertical stripes, indicating that they were not specific to movement direction. **(F)** Contributions from three sets of extrinsic input. In the panels where the input was modulated in the corresponding epoch (early-early, middle-middle, and late-late), there was a prominent diagonal banding, showing that these contributions could impart directionality to the firing rate of the output neuron.

#### 2.2.1 Recurrent HNN

We consider the neural activity recorded from primary and premotor cortices to be a sample from a large, interconnected network of neurons involved in generating behavioral output. With this consideration, we modeled the activity during our experiment by embedding this sample as a subset of neurons in the hidden recurrent layer of an artificial neural network. The recurrent layer acts as a local network around the neurons whose activity we recorded empirically. All neurons in this layer receive connections both from intrinsic neurons and from those that are extrinsic, as described in the previous paragraph. This allows us to distinguish the effect of each type of input on the modeled version of our recorded activity.

Recurrent neural networks can be trained to produce temporal patterns of output using only internal feedback pathways of different duration. Such a model was developed by Sussillo et al. [22], to show how recurrent neurons, provided with *transient directional*

*input* to set an initial condition, could produce subsequent temporal patterns of muscle activity during reaching. We used an RNN with a similar architecture to instead show how *ongoing extrinsic input* can interact with recurrent activity to generate models of the firing rates we observe in our data (Methods and Fig.2A). After training, we found that the firing rates of neurons in the hidden layer were cosine-tuned, that the models of the empirical firing rates were highly accurate (mean correlation = 0.91, Monkey C, 0.96 for Monkey N; Fig.2B), and that the overall population underwent the same state transitions as those in our data (Fig.2C).

The interplay between intrinsic and extrinsic factors can be described using the correlational structure of the RNN activations, **x**(*t*) (Eq.2), as they evolve during the movement [23]. To visualize the dynamics of these correlation patterns, we carried out rPCA on the activations and plotted their projections on three of the axes to form neural trajectories (Figs.2D, S5C, Methods). Since the profiles of these scores overlap in time, the resulting neural trajectory, when plotted in a 3D manifold, is curved. 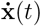 is formulated as a linear combination of input from recurrent connections and external inputs, so that the tangent along the trajectory can be divided into components that are intrinsic (purple arrows) and extrinsic (green arrows). While the intrinsic dynamics due to recurrent connections always tend to pull neural trajectories toward the center (a point attractor), the external inputs drive individual neural trajectories in different directions (green arrows) during individual epochs of the movement. Each trajectory segment in these different directions corresponds to one of the network states (gray planes in Fig.2D) and shows that the extrinsic input is driving the changes in dynamics of the model. The same results were found for data collected from Monkey N (Fig. S5).

##### 2.2.1.1 Input Contributions

The defined connectivity of the trained RNN, combined with the prominent tuning of the neurons in the network, make it possible to investigate how inputs are combined to produce the firing rate patterns of the individual neurons in our data set. To understand how directionality is imparted to the output neurons serving as models of our empirical results, we analyzed the differences between the effect of contributions that are intrinsic (recurrent layer) and extrinsic (input groups) (Eqs.4, 5) to the modeled firing rates. The contributions to the firing of RNN-59C are visualized in separate heat maps for the intrinsic (Fig.2E) and external (Fig.2F) inputs in each epoch (see Methods and Fig. S5 for another example unit). Since both input sets had cosine-tuned firing rates (prescribed for the extrinsic units, calculated for those in the recurrent layer), they were grouped by preferred direction (heat map columns) and evaluated in each movement direction (heat map rows).

The preferred direction of the input in each column coincides with the direction of movement along the diagonals of the heatmap. As expected, this is where we see the largest positive and most negative contributions for movement toward the preferred and anti-preferred directions, respectively. This expectation holds for the extrinsic inputs (Fig.2F), but not for those that are intrinsic. In the early epoch (bottom panel Fig.2E), the positive and negative contributions are reversed, which, by itself, would lead to output tuning opposite to what was observed. The intrinsic heatmaps for the other epochs have vertical bands which are positive for input that matches the output preferred direction and negative bands in the anti-preferred direction. Since there is a similar positive and negative contribution of input for each movement direction, these would tend to cancel out, again showing that the intrinsic input cannot account for output directionality.

Strong contribution along the diagonal promotes directional tuning in the output unit. These results and those of two additional analyses, Input Correlation (Fig. S8) and Input Dropout (Fig. S9), show that the extrinsic input was the dominant factor governing the directional tuning in the RNN model across all units in our data.

#### 2.2.2 Spiking neural network

Motor cortical activity is a function of both intrinsic and extrinsic connectivity. Nonetheless, based on the results from the RNN, we asked whether *extrinsic* input, by itself, could produce the temporal patterns of firing we observed in our data. Again, we assumed that the three profiles uncovered by rPCA (Fig.1D) signified common input and modeled this as three groups of 90 tuned neurons modulated at different points in the reach. Each neuron’s firing was driven by a Gaussian temporal profile. In addition to these three sets of directional input, a fourth group was used to impart a non-directional speed offset [24, 25] to the firing rates (Fig. S12A). This group was driven by the movement speed profile shifted 50 ms backward, with gains that varied from −0.5 to 1, across the 90 neurons.

For an individual trial, the drive to each input was transformed to spike trains (visualized as firing rates in Fig.3A) using the integrate-and-fire algorithm in the Brian2 simulator [26]. On individual trials (repetitions 1-20) during training, we applied Hebbian learning using the spike occurrence times of the recorded units and those of all input units to find input-output weights (Figs.3B, S12B). After training, modeled firing rates of the output neurons were generated with the SNN by aligning the input groups to the behavioral landmarks during the trials (21-40) not used for training (Figs.3C, S7). Firing rates of the recorded neurons were well fit by the modeled output. (r = 0.71 across all units for Monkey C and r = 0.69 for Monkey N). However, two factors led to reduced SNN performance compared to the RNN: the SNN struggled to capture shifts in baseline firing rates, and it performed poorly when the recorded unit’s modulation did not align with the time course of input unit modulation. Population dynamics of the modeled data matched that of the actual data (compare Fig.5A to Fig.1D), showing that the model captured the network state transitions. The common input, by itself, drives both the predicted firing rates and the state transitions of the network.

**Figure 3.**
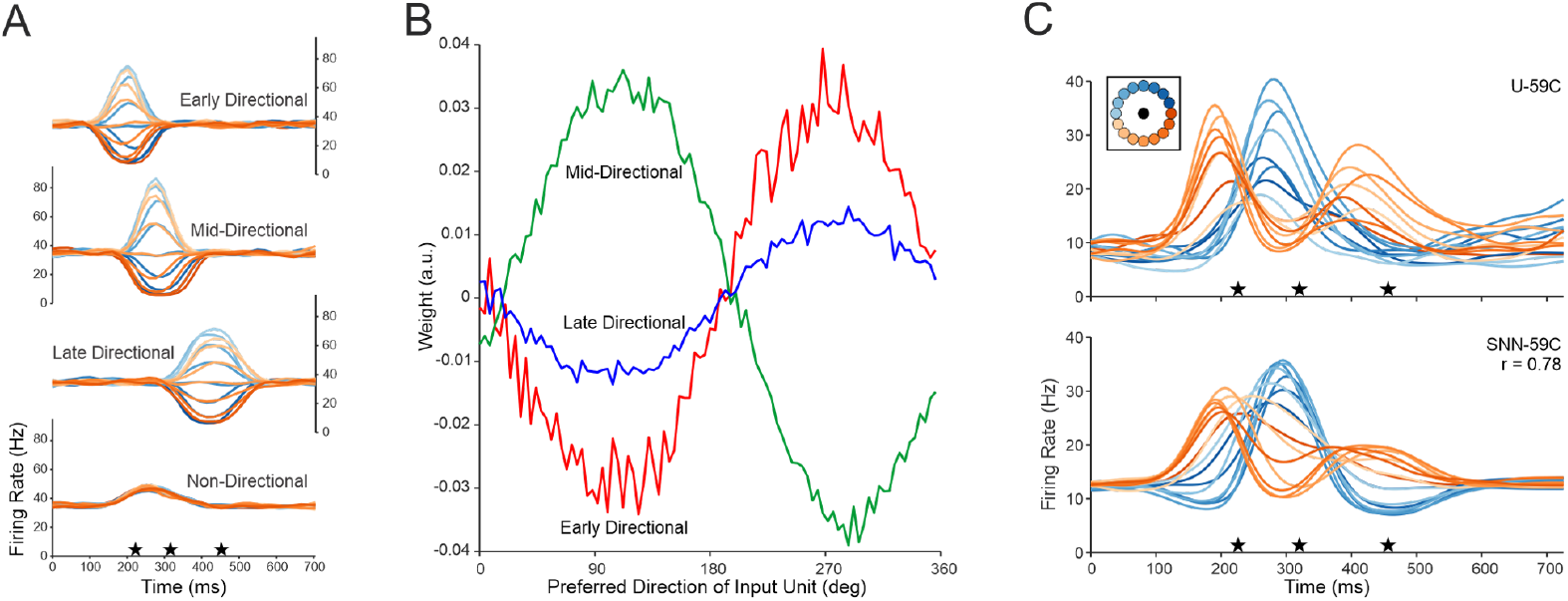
Spiking neural network. **(A)** Trial-averaged firing rates for one example input neuron in each group. The three example neurons were tuned to 180°. Stars indicate average time for movement onset, peak speed, and movement completion. The non-directional example neuron had an amplitude coefficient of 0.25. **(B)** Weights for the directional input units to modeled unit (SNN-U59C). **(C)** Actual and predicted modeled firing rates for Unit 59. The stars mark the behavioral events as in **A**.

##### 2.2.2.1 Statistical structure of synaptic integration

The SNN model explicitly captures the postsynaptic effects from presynaptic connections. Using a single example neuron (SNN-59C), we used spike-triggered averaging [27] to show a gradual rise in membrane potential 20 ms before an output spike, followed by a sharp increase just before spiking (Fig.4A). Accordingly, we considered synaptic integration in two intervals before an output spike: *buildup* – 20 ms (Figs.4 C, E) and *trigger* – 0.1 ms (Figs.4D, F). These analyses were carried out separately in distinct portions of the reach (Fig.4B) corresponding to the three epochs of modulation.

**Figure 4.**
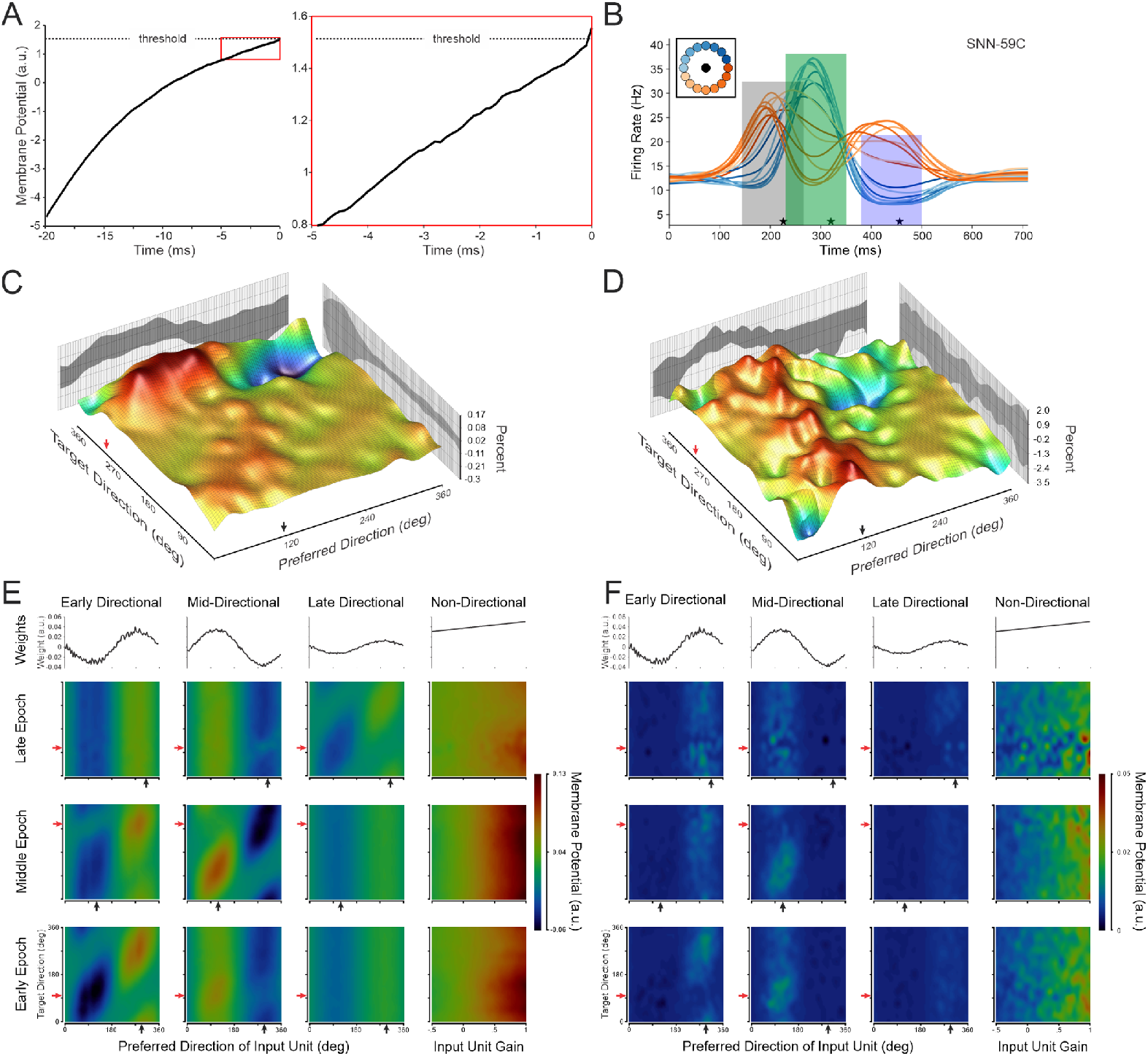
Input contributions to output spiking. Exemplified with SNN-59C. **(A)** Spike-triggered average of membrane potential. The membrane potential of the modeled unit before an action potential was averaged across its spikes in the middle epoch time zone (green rectangle in **B**) for a reach at 112 degrees. (Left) 20 ms average before the spike. (Right) 5 ms before the spike. **(B)** Analysis zones for the three epochs defined by the rPCA analysis. The center of each zone was determined by the peaks of the rPCA components and then shifted on each trial relative to the three behavioral landmarks (stars). **(C)** Buildup-Percent of spikes in the buildup window for each input neuron in the mid-directional input group during the middle epoch in each movement direction. The chance percent of a spike in the window has been subtracted out. **(D)** Percent of trigger input spikes. Same as C, except the window is now 0.1 ms. The back panels of C and D show the projected input unit percentages. The side panels show the projected percentages across movement directions. **(E-F)** Heat maps of the change in the membrane potential of SNN-59C during the buildup **(E)** or for the trigger input **(F)**. Columns correspond to the input groups, the weights for each input group are shown in the top row, and the other rows correspond to the epochs shown in **B.** Black arrows indicate the epoch-specific preferred direction of the SNN-59C; red arrows are in the corresponding anti-preferred direction.

**Figure 5.**
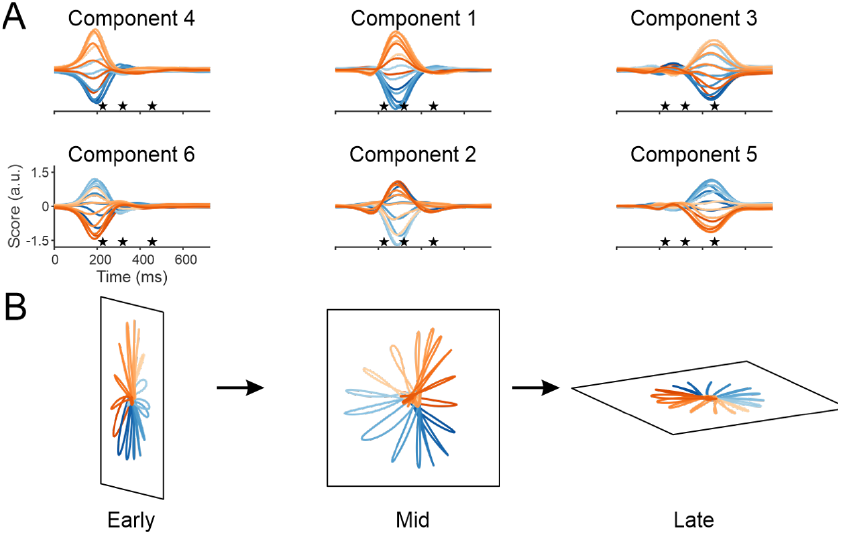
Discrete dynamic components (data from the SNN-predicted firing rates-Monkey C). **(A)** Three pairs of rPCA scores defined by coincidence of their peak values. The stars denote the mean behavioral landmarks of movement onset, peak speed, and the end of movement. The traces begin when the target was presented. **(B)** Each of the three pairs was plotted on separate axes of a plane to form a manifold showing how the correlated structure in an epoch varies during movements in different directions. The orientation of the three planes was chosen to emphasize the state transitions associated with each epoch. Movement direction is color-coded.

In the SNN, output spiking is influenced by the input neurons’ firing rate and input-output weight. The number of input spikes before an output spike is a function of the covariation of input and output firing rates and of the relative timing between input and output spikes, which has been defined as “functional coupling” [28]. We found functional coupling by removing the contributions of firing-rate covariation (Methods). The probability of observing functional coupling increased for input neurons with tuning near the output neuron’s preferred direction in the buildup phase, with an especially strong effect for reaches toward the anti-preferred direction (Fig.4C). In contrast, the probability of functionally coupled triggering increases for input neurons with large positive weights, regardless of movement direction (Fig.4D).

Spike timing alone does not completely describe how inputs are combined to generate output spikes. Synaptic integration in the SNN can be understood by describing how the output unit’s *membrane potential* develops before an action potential. The change in membrane potential was calculated using spike timing and network weights, for each input. Input neurons fired most rapidly when their preferred directions aligned with the reach direction, forming a diagonal pattern on the heatmap during the buildup (Fig.4E). Despite varied contributions, the SNN input-output pattern was like that found for the extrinsic input to the RNN (Fig.2F), suggesting consistent input-output patterns across architectures. The diagonal pattern of contributions in the SNN heatmap was evident across the activity in the data sets from both monkeys (Diagonal Test – Methods).

The input neurons acting as *triggers* (Fig.4F) are those with the largest weights. This weight-dependency is readily apparent for the late epoch (Fig.4F-top row). Even though the neurons in the late input group fire much faster in this epoch than those in the other groups, their relatively small weights (weak connectivity) make them unlikely to trigger an output spike. This was a consistent finding for our entire data set (Fig. S11).

Non-directional inputs, aligned with reach speed profiles, contributed primarily during the early and middle epochs, with a large effect on neurons with single-epoch firing rate modulation (Unit SNN-48N in Figs. S7 and S10). Whereas these inputs tend to modulate the firing rate of neurons in the middle of the reach, the directional tuning of each output unit comes from the three sequential sets of directionally tuned input neurons. Neurons with preferred directions matching the movement direction contribute to the buildup membrane potential toward threshold, while those with the largest weights (tuning matching that of the output neuron) trigger the spike.

## 3 Discussion

Using our two HNN models, we found that extrinsic input, **u**(*t*) (Eq.1) is a major explanatory factor of our empirical results. We first built a recurrent neural network (RNN) to examine the relative effects of *intrinsic* and *extrinsic* input on the firing rates of the neurons we recorded empirically. This model showed that the extrinsic input was responsible for generating the directionality of the recorded neurons. Based on this result, we asked whether a simple feedforward spiking neural network (SNN) driven *only* by extrinsic input could account for the empirical activity in our recorded data. The SNN accurately modeled the firing rates of the recorded neurons as well as the individual temporal patterns of correlation in the overall population, supporting the role of continuous input to motor cortical areas during reaching.

We found that correlations between the firing of neurons, detected through dimensionality reduction, arise largely from extrinsic inputs rather than intrinsic connectivity. While firing-rate correlation between a pair of neurons might imply that they are connected [22], given the inter-electrode spacing of the Utah probe, it is more likely that the correlations came from indirect connections [29, 30]. Multiple studies have shown that polysynaptic and shared input can drive these patterns [21, 31–37]. In our models, these correlations reflect directional tuning as a global property of network function [20, 29]. During a straight reach, state transitions are driven by successive groups of input neurons modulated across time. Input neurons with similar tuning across groups fire rapidly for movements in their common preferred direction, but their impact varies because corresponding neurons in each group have different input-output weights. This reflects the concept of latent input: anatomical projections to an output neuron may be hardwired, but they will not be connected functionally unless they are active. Our results suggest that this functionality is determined by behavior events, such as movement onset, peak speed, and target acquisition, which align with distinct network states. Reaching is divided classically into a “ballistic” and “homing phase” [38], but more recent studies show that, although the first portion of the reach is rapid, it is also modifiable by ongoing extrinsic input [39, 40]. Our results are consistent with these behavioral results. Furthermore, we have shown that the late epoch activity is diminished when visual accuracy requirements are removed [12, 41]. Together, these observations show that ongoing behavior corresponds to the latent input, driving cortical state transitions.

We considered two causal components of spike generation using the SNN: buildup and triggering. Contributions to membrane potential during buildup were governed both by the firing rates and weights of the input units, while those that triggered the output spikes were determined primarily from the weights of the input, regardless of firing rate. Input neurons contributing to the membrane buildup had a broad range of preferred directions, while those that triggered the spike were tuned to a narrow range around the output neuron’s preferred direction. This finding is consistent with those from an *in vivo* mouse study in which the membrane potentials of visual cortical neurons were recorded as oriented visual input was presented [42]. A wide range of orientations increased the membrane potential, but only those matching the output orientation triggered a spike. The buildup and trigger phases of spike generation have been considered in a computational model showing that combined excitatory and inhibitory input contributions (during buildup) can set the “working point” [43] of an output neuron to a critical range, making it sensitive to additional input (triggers) that can change its effective connectivity [28, 32]. Variations in the composition of the contributions during buildup can explain why ineffective inputs during one state can become effective in another.

Here we are focused on the disparate *input* driving the firing rates we observed. Although we built the HNNs with different groups of episodic extrinsic input, an alternative possibility is that the weights of a single group could change on a rapid timescale through molecular synaptic mechanisms [44]. The SNN results show that extrinsic drive, by itself, can produce the firing rates we observed in our data. However, more realistic spiking neural networks can be expected to show that intrinsic recurrent input has an important role in network functionality.

Using these models, we can address two issues in the current literature, both stemming from the assumption that there is only a single process taking place during reaching. Using the concept of dynamical systems (Eq.1), a set of studies (reviewed in [45]) concluded that the change of firing rate, 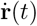, is a function of neuronal firing rates operating only through intrinsic connectivity (i.e.,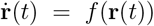) during a reach. This conclusion was based on the observation of PCA components that formed curved neural trajectories. These trajectories were taken as “rotational dynamics” and interpreted as a signature of a single, first-order dynamical process. Our findings show that the curved trajectories are formed from pairs of overlapping temporal components. By removing the overlap, our neural trajectories were straight. Using rPCA, we identified distinct pairs of contemporaneous latent variables that appear at consistent points in the reach. The pairs of latent variables from the SNN simulations closely matched those of the empirical data (Figs.1D, 5A-B). When the sequential pairs of components are plotted on a sequence of manifolds, they form straight neural trajectories (Fig.5), consistent with a series of separate cortical states during a reach. Using our methodology, this shows that instead of a single process, cortical activity evolves through separate discrete network states associated with different components of the reach.

Our finding of distinct behavior-related neural states also provides an alternative to the conclusion of a recent study in which monkeys performed a reaching task by visualizing a neural trajectory of motor cortical population activity [46]. Monkeys were able to perform the task in the forward direction by starting at the beginning of the neural trajectory but failed when they needed to reverse the trajectory. This was taken as a sign that intrinsic “dynamical structure is highly conserved and not readily changeable.” Our results showing that the dynamical structure is driven by extrinsic, behaviorally linked input provide a different explanation: the reach termination state cannot precede reach initiation.

Complex functioning of neural systems is now beginning to be described using network models. Recognizing patterns of neuronal activity within a neural population is a first step in characterizing the complexity of network functionality [28, 47]. In realistic networks, the concepts of input, output, and modularity are ill-posed [48]. For instance, the idea that arm direction is the only input to a cortical neuron during a reach is overly simplistic, as tuning functions encompass many parameters simultaneously [49–51]. Defining the boundary of the generative network (for instance, by constraining it to the motor cortical areas) is unrealistic. Even the current designations of intrinsic connectivity and extrinsic input are abstract constructs. Although it is generally agreed that state transitions in biological neural networks are synaptic phenomena [19, 52–56], the organization of events leading to a state transition is an open question. Even though our models do not directly address the mechanical details underlying behavior generation [57], the observed sequential patterns of neuronal correlation can be viewed as signatures of separate functional operations occurring in a network that includes, but is not limited to, those in motor cortical areas. Consistent with the idea of distributed systems, the same population of neurons participates in each of these neural operations. The simple SNNs in our study can be enhanced by adding biological properties of synapses such as channel conductance, transmitter species, receptor properties, and structural anatomy. Local intrinsic connectivity can be studied by including a hidden layer in the SNN. New experiments that combine large-scale recordings with direct observation of membrane potentials can be used to verify and extend these models. Models that combine artificial and empirical neural activity make it possible to preserve the complexity of realistic neural systems which will lead to plausible operational principles responsible for the generation of behavior.

## 4 Methods

### 4.1 Behavioral task

Two male rhesus macaques (monkeys C and N) performed a center-out reaching task using a virtual reality setup. Each monkey sat in a primate chair with one arm restrained. An infrared marker was placed on the wrist of the freely moving arm. The position of the marker was sampled at 60 Hz (Monkey C) or 100 Hz (Monkey N) (Optotrak, Northern Digital) and projected as a spherical cursor on a 3D monitor (Dimension Technologies). At the start of each trial, a spherical target appeared in the center of the virtual space and the monkey was required to hold the cursor within the target for 200-300 ms. The center target was then extinguished and a peripheral reach target (radius of 10 mm for Monkey C, 6 mm for N) presented at a position randomly selected from 16 predefined locations. The 16 peripheral targets were radially arranged around the center target on a 2D virtual plane in front of the monkey (target-to-center distance of 7.4 cm for Monkey C,6.5 cm for N). Trajectories and speed profiles of the reaching movements are shown in Supplementary Fig. S1. A liquid reward was administered after the monkey reached and held within the 3D target sphere for 200-400 ms. All procedures were in accordance with the guidelines of the US National Institutes of Health and were approved by the Institutional Animal Care and Use Committee of the University of Pittsburgh.

### 4.2 Neural recording

Monkey C was implanted with one multi-electrode array (96 channels, Blackrock Microsystems) in the arm area of the primary motor cortex contralateral to the freely moving arm, and Monkey N had two arrays in the dorsal premotor cortex (Supplementary Fig. S2). Single-neuron responses were isolated using the Offline Sorter program (Plexon Inc.). Ninety-three single units were recorded from Monkey C, and 113 units from Monkey N. The Monkey C data have been used in previous studies [12, 25], and the Monkey N data are novel.

### 4.3 Data pre-processing

To perform trial-averaging, center-out reaching trials were aligned at a series of behavioral landmarks,including peripheral target presentation, movement onset, maximum speed, movement offset, and target hold off. Movement onset and offset were defined as the times when speed reached 20% of the maximum. Trial alignment was implemented by fixing the number of time bins in the epoch between a pair of adjacent landmarks for all trials. The number of time bins in an epoch was determined by the average duration of that epoch across all trials divided by bin size (e.g. 5 ms per bin). The number of fractional interspike intervals in each bin was divided by the bin size to get firing rate [58, 59]. The spike rates were smoothed with a Gaussian kernel (25 ms SD). For each neuron, firing rates were then averaged across trials with the same target (47 repetitions of each target for Monkey C, and 49 for N). The mean firing rate, over time and across all task conditions, was calculated for each neuron. Neurons with low mean firing rates (*<* 1.5 spikes/s) were excluded from subsequent analyses, leaving 67 units for Monkey C and 78 for Monkey N. A “soft” normalization” [60] was used to normalize firing rates, so that all neurons had similar ranges after the process. Hand position was spline-interpolated and then resampled to match the number of bins in each epoch. Velocity was then recalculated using resampled positions. All data processing and subsequent data analyses were performed using customized Matlab (Mathworks Inc.) scripts, except that the spiking neural network modeling (see below) was performed with Python codes.

### 4.4 Principal component analysis with rotation

Each trial-averaged firing rate data set was combined into a matrix of size *c* × *t* × *n*, where *c* is the number of conditions, i.e., movement directions, *t* is the number of time bins, and *n* is the number of neurons. Non-directional components were first calculated for single neurons by averaging the *c* firing rate profiles across conditions. These non-directional components were then subtracted from the profiles of each neuron to yield the directional components.

The directional and non-directional modulation of the recorded units were analyzed following the general approach of Suway et al., [12] – the directional component was fit to a cosine function for each time bin and each unit, which uncovers the amplitude and fit (R^2^) for directional modulation. One through four Gaussians were fit to this directional modulation profile. The number of Gaussians was chosen to fit the data well, and to cover the areas of data that had good cosine fits – with a penalty to discourage using additional Gaussians without explaining significantly more variance. We tested for the presence of modulation in the non-directional component by determining if the profile changed at least 1.5Hz over the course of the trial.

Principal component analysis (PCA) was used to reduce the dimensionality of data. The previous firing rate data matrix of size *c* × *t* × *n* (with non-directional components removed) was reorganized to be matrix *X* of size *ct* × *n*, by concatenating firing rate profiles of all conditions for individual neurons. Each row of *X* was a 1× *n* vector of firing rates from the neuronal ensemble, and there were *ct* samples from all time bins and conditions. PCA found a set of *n* orthogonal axes, i.e., principal components (PCs), in the *n*-dimensional firing rate space, such that the top *k* (*k*≪ *n*) PCs defined a subspace that explained the most variance of the data. The data matrix *X* was accordingly decomposed as:

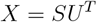

where matrix *U* of size *n* × *n* had *n* eigenvectors as columns, *U*^*T*^ was the transpose of *U*, and matrix *S* of size *ct* × *n* contained PC scores. Each row of *S* was a 1× *n* vector of scores, which were projections of the corresponding row of *X* onto the *n* eigenvectors. Columns of *U* were ordered by the amount of data variance explained by PCs, from high to low. By only keeping the top *k* PCs, we reduced the dimensionality of *X* from *ct* × *n* to *ct* × *k*, and had

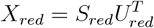

where *X*_*red*_ of size *ct* × *n* contained reconstructed firing rates using top PCs and scores, *U*_*red*_ of size *n* × *k* had the first *k* columns of *U*, and *S*_*red*_ of size *ct* × *k* had the first *k* columns of *S*. In our analyses, we chose *k* to be 6, thereby accounting for 80% of the variance of firing rates in the population for Monkey C (86% for Monkey N).

To reinforce simple structure and improve interpretability, Promax rotation was applied on the *S*_*red*_ matrix of PC scores. A target matrix using the Equamax method was found first. A transformation matrix, *R*, of size *k* × *k*, was then calculated such that *S*_*red*_*R* conformed best to the target matrix with respect to the sum of squared errors criterion. *X*_*red*_ can be decomposed as

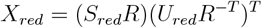

where *R*^−*T*^ was the transpose of the inverse matrix of *R. U*_*red*_*R*^−*T*^ was a matrix of size *n* × *k* and its columns were the rotated eigenvectors (rPCs). *S*_*red*_*R* of size *ct* × *k* consisted of rotated scores (rPCA scores). The transformation matrix *R* was found using the Matlab *rotatefactors* function. Note that the rotated axes were not principal axes (or principal components) anymore, and they were no longer required to be orthogonal to each other. We refer to this PCA with rotation approach as rPCA.

We illustrate the difference between rPCA and standard PCA, using a toy dataset containing the firing rate profiles for reaching movements in 16 directions from 3 simulated neurons (Supplementary Fig. S3A). The firing rate of each neuron is directionally tuned during 3 temporal epochs centered at 0.3 s, 0.45 s, and 0.7 s, respectively. Around the center of each of the 3 epochs, a firing rate modulation “hump” (has the shape of the probability density function of a Gaussian distribution) was added to the constant baseline firing rate of each neuron. The amplitude of each hump equals the modulation depth (a scalar) multiplied by the cosine of the angle between the movement direction and the neuron’s preferred direction during that epoch. The modulation depth for each neuron in each epoch was randomly chosen from a uniform distribution between 0 and 1. The preferred directions of all neurons are either the same or 180 degrees apart during the same epoch, such that the neural trajectories during those periods of time follow straight lines (Supplementary Fig. S3D, seg. 2, 4 and 6). The preferred direction changed randomly between epochs. The width of the firing rate humps was determined by the standard deviation of the Gaussian distribution, which was chosen to be 0.04 s, 0.05 s and 0.09 s, respectively, for the three epochs. This created an overlap between two adjacent epochs, and therefore the neural trajectories tend to curve during these overlaps (Supplementary Fig. S3D, seg. 3 and 5). The standard PCA could not isolate those directionally tuned modulation epochs from one another, since each PCA component was active in all three epochs (Supplementary Fig. S3B). In contrast, the rPCA successfully separated those epochs (Supplementary Fig. S3C) and the rPCA axes (black straight lines in Supplementary Fig. S3D) are well aligned with data samples from the nonoverlapping portions of those epochs. Therefore, PCA with rotation was used as the default dimensionality reduction method in this study.

### 4.5 Recurrent Neural Network (RNN)

We trained recurrent neural networks to replicate neural activity recorded from the primary motor cortex during reaching movements. Each of the *N* (=1,000) units in the RNN behaves following a differential equation of the form

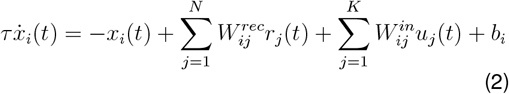

where *x*_*i*_ denotes the activation of unit *i* and *r*_*i*_ its firing rate; *u*_*j*_ represents the *j*-th extrinsic input; 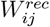 is the recurrent connection weight from unit *j* to *i*, and 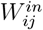 the weight from input *j* to unit *i*. Each unit has a bias, *b*_*i*_. A single time constant *τ* is used for all units in the network.

Extrinsic inputs, *u*(*t*), were created based on the rPCA components of the recorded motor cortical neural activity during reaching. There were three groups of extrinsic inputs, corresponding to the three pairs of rPCA components (e.g. Fig.1D). Each rPCA component was fit with a Gaussian kernel to find the timing of the peak (*µ*_*i*_, *i* = 1, 2, 3) and the width (represented by its standard deviation *σ*_*i*_). The temporal offsets between peaks of rPCA components and the three behavioral landmarks were then calculated as Δ*t*_*i*_ = *µ*_*i*_ *™ t*_*i*_, where *t*_*i*_,*i* = 1, 2, 3 represent trial-averaged timing of movement onset, maximum speed, and movement offset respectively. Note that rPCA analysis and the above behavioral landmarks (*t*_*i*_) were based on data temporally aligned across trials (see Data pre-processing subsection). When creating extrinsic inputs for RNN in individual trials, the first extrinsic input group followed

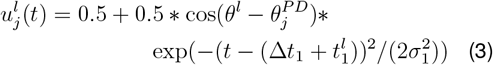

where *θ*^*l*^ represents target direction in trial 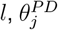 is the preferred direction of the *j*-th input (*j* = 1, 2, …, 18), and 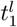 is the time of movement onset in trial *l*. The remaining two input groups were similarly defined using timing of maximum speed and movement offset respectively. There are 18 inputs in each of the three input groups (therefore *K*=54), with preferred directions distributed evenly around a circle at 20-degree intervals. For the input dropout analysis (Supplementary Fig. S9), the latter two groups of inputs were set to a constant value of 0.5 for all times. This demonstrates how RNN outputs would change after losing some of the extrinsic inputs.

The activation, *x*_*i*_(*t*), of a recurrent unit was converted to firing rate via the rectified hyperbolic tangent function,

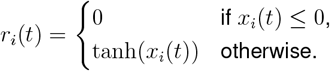

The RNN has *M* output units, which are made identical to a subset of *M* randomly selected recurrent units of the network (Fig.2A, nodes in blue). Networks is were optimized such that the activities of the M output units match firing rates of recorded motor cortical neurons. Training of RNNs aimed to minimize the cost function

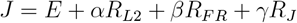

where *E* is the squared error between network output and recorded neuronal activity; *R*_*L*2_ is a standard L2 penalty on the input weights and recurrent weights; *R*_*F R*_ helps prevent permanent saturation of RNN units; *R*_*J*_ encourages simple state-space trajectories. Hyperparameters *α* (=1e-5), *β* (=1e-3) and *γ* (=1e-5) are weights of regularization terms. Detailed explanation of the RNN model can be found in Sussillo et al. 2015 [22].

All single-trial data were randomly divided into 5 folds. The RNN model was trained with 4 folds of data and then tested on the remaining fold. This was repeated 5 times such that each fold of data acted as the test dataset once, i.e., 5-fold cross validation. Training of RNN models was performed using Bridges-2 [61] at Pittsburgh Supercomputing Center.

#### 4.5.1 Extrinsic and Intrinsic Contributions to RNN Units

After the RNN was trained, we calculated the contributions to the change of activation of individual RNN units. According to Eq.2, contributions to 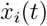 come from two sources, the external inputs and the intrinsic recurrent network structure. The mean contribution, during a temporal epoch between *t*_1_ and *t*_2_, from external input *j* to RNN unit *i* was calculated following Eq.4.

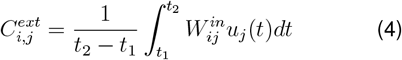

Repeating this analysis for all 16 movement directions and all 54 external inputs yielded a 16 ×54 matrix. This matrix was then split into three 16× 18 matrices, each corresponding to one of the three extrinsic input groups. These matrices were visualized using heatmaps and are shown as one row of the right panel in Fig. 2E. This analysis was performed in three temporal epochs (early, middle, and late) during the trial. Each epoch was 120 ms in duration, with its midpoint aligned to the peak of the corresponding group of bell-shaped inputs (see Eq. 3). These same epochs were also used when analyzing the SNN model (see Fig. 4B).

To compute the contributions from intrinsic network structure, all RNN units were divided into 18 groups according to their preferred directions during the epoch. The combined contribution from the *k*th RNN unit group, *G*_*k*_, was calculated as using Eq.5

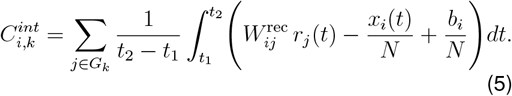

Results of the 18 RNN unit groups for 16 movement directions during each of the three temporal epochs were presented as heatmaps in Fig.2E left panel in a similar way as described above.

#### 4.5.2 Directionality Test

The differential extrinsic and intrinsic connections contributions to the directional tuning of an RNN unit, were calculated using the correlation coefficients between tuning curve of the RNN unit and those of extrinsic and intrinsic contributions, respectively.

To get the tuning curve of the intrinsic contributions, the 16 ×18 heatmap explained above for a given temporal epoch was added up across all columns, resulting in a 16× 1 vector representing the total contributions for the 16 movement directions from all RNN units. Similarly, the tuning curve for extrinsic inputs, a second 16× 1 vector, was found by adding all 54 columns of the three 16× 18 matrices of contributions from the three input groups. The mean firing rate of the target RNN unit in that epoch was also calculated for all movement directions, forming a third 16× 1 vector. The correlation coefficient between the vector of contributions and the firing rate vector was then calculated, for extrinsic and intrinsic contributions, respectively. This analysis was repeated for all three temporal epochs and for all RNN output units. The distribution of these correlation coefficients across all output units in each epoch is shown as a histogram in Supplementary Fig. S8. This analysis reveals that contributions from external inputs had stronger correlation with output firing rates than those from intrinsic drivers.

#### 4.5.3 RNN State Space Trajectory

The state-space trajectories of the model were visualized by performing rPCA on the activations of all RNN units, **x**(*t*) = [*x*_1_(*t*), …, *x*_*N*_ (*t*)]^*T*^. A 3-dimensional state space was formed using 3 of the top 6 rPC scores, one from each of the three pairs of components. Single-trial **x**(*t*) was then projected into this 3-D space to find the visualization of state-space trajectory (Fig.2D). The gradient, 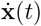, was then calculated along the trajectory. Similar to the contribution analysis above, we split the gradient into two parts as driven by extrinsic inputs and intrinsic network structure, respectively. These two parts of 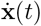 were then projected back to the above 3-D space and plotted as arrows in Fig.2D (green arrows as from extrinsic inputs and purple for intrinsic dynamics). A 2-D flow field within this 3-D space was made to further visualize the intrinsic RNN dynamics. Evenly spaced sample points ([*s*_1_, *s*_2_] value pairs) were taken from this 2-D plane. The averaged values across time and trials of the remaining *N™* 2 dimensions were calculated (a vector of [*s*_3_, …, *s*_*N*_]), and then combined with each of the above 2-D sample points respectively to form *N* -D vectors. These vectors were then projected back to the original *N* - D space using rPCA axes to find the corresponding **x** = [*x*_1_, …, *x*_*N*_]^*T*^ vectors. Using Eq.2 above, we calculated 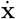 for each **x**, and then split it into two parts, 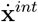 and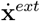, as due to intrinsic network structure and extrinsic inputs, respectively. The 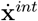 vector for each sample point was projected back to the previous 2-D plane and plotted as a purple arrow for each sample point on the bottom plane of Fig.2D.

### 4.6 Spiking Neural Network (SNN)

The Spiking Neural Network (SNN) was constructed to replicate recorded neural activity from a set of simulated extrinsic inputs. The network was constructed using four groups of 90 neurons which were fully connected to the output neurons (Supplementary Fig. S12A). Following the same motivation as that for the RNN, the network was trained so that the firing rates of the output neurons matched those of the neurons recorded empirically during reaching.

The input comprised three sets of 90 neurons with directional tuning during the epochs identified by rPCA, and one non-directional set of 90 neurons tuned to reach speed. The Brian2 SNN simulator [26] was used to generate input spikes from an underlying drive function. For the directional inputs, the drive was modeled as a Gaussian fit to the modulation of the rPCA data (as described above in the RNN section), scaled by the cosine of each neuron’s preferred direction, which was evenly distributed around a circle. These profiles were further scaled by movement speed and included a positive offset to increase the baseline firing rate, with additional noise to reduce correlated firing between input units. The peaks of the profiles in each input group were aligned to the behavioral landmark in its corresponding epoch in each trial. For the non-directional group, the drive followed the hand’s speed profile, shifted backward by 50 ms, with neurons having a gain distributed from –0.5 to 1.0, plus an offset and noise.

The weights were trained using the relative timing of input and output (recorded) neuron spikes. The potential contribution to weights for each input neuron was tracked individually. The baseline contribution is negative and at the time of each spike, a two-sided decaying exponential was added onto the baseline. The value of the potential contribution was added to the weight between the input and output unit at the time of the output unit spiking. If input and output neuron’s spikes coincided, a positive contribution was added to the weights, otherwise a negative contribution was added. The baseline contribution was adjusted so the average weight into an output unit from an input group was zero (Supplementary Fig. S12B).

The network was optimized by scaling the weights from each input group into each output and by adjusting the modeled output unit thresholds to minimize the RMSE between predicted and actual firing rates. A differential evolution optimization method [62], was used to iteratively evaluate parameter combinations and combine the best performers for the subsequent iteration evaluation.

Finally, the network was simulated for all trials using the optimized weights and thresholds. The membrane potential for each output unit was tracked, with changes occurring whenever an input unit fired, influenced by the connecting weight. The membrane potential also tended towards a resting potential.

#### 4.6.1 Statistical structure of synaptic integration

Analyses were performed to explore the statistical structure of input spikes leading to output spikes in the SNN model. Because of the deterministic quality of the SNN model, the precise effect on output neuron membrane potential from of each input spike can be quantified. To find which input spikes lead to an output spike during each of the three epochs, we restricted analyses to three distinct segments of the reach (Fig.4B), as described in the main text. Input spikes occurring within a 20 ms (*build-up*) or 0.1 ms (*trigger*) window before output spikes in these segments were considered for analysis.

First, raw spike counts were extracted. There were 360 individual input units (90 units × 4 input groups), 16 target directions, and 3 epoch segments (Fig.4E-F). Input spikes occurring in either window before an output spike were counted. To get spike probability, the spike counts within each target/epoch combination were divided by the total number of input spikes occurring in the target/epoch. Finally, to get the weighted contribution to membrane potential as shown in Fig.4E-F, the spike probability was multiplied by the input’s corresponding weight. These analyses were completed for both the *build-up* and *trigger* windows.

We also wanted to interrogate the importance of the relative timing of input spikes in causing output spikes. To do so, we determined the spike probability that would be expected by chance. The random occurrence of a spike in the window was calculated by redistributing the spike times of all input and output units within each epoch segment for all trials. The spikes that occurred in each trial and epoch segment were reassigned a new location within the epoch based on a random uniform sampling of locations within the epoch. Afterward, raw spike counts and spike probabilities were calculated as described above. To generate the surfaces seen in Fig.4C-D, the spike probability expected by chance was subtracted from the spike probability generated from the actual spike times.

Additional analysis was done with the weighted contribution. Fig.4E shows a strong diagonal, which indicates input units with a preferred direction that matches the direction of movement have relatively large contributions to the output unit membrane potential. We quantified this phenomenon using the Directionality Test (below) for all units in our data set.

#### 4.6.2 Diagonal Test

We down-sampled the 90 columns of the heatmaps to 16 in order to create a square (16× 16) matrix and then calculated a diagonality metric, which is simply the sum of the magnitudes of contributions along the diagonal divided by the sum of all 256 contribution magnitudes. This metric was calculated for all output units, epoch segments, and the three directional input groups for the *build-up* window. A bootstrap analysis was performed on each matrix by randomly selecting 16 of 256 contributions in the matrix (without resampling each row or column) and calculating the diagonality metric using these contributions as the “diagonal”. This was repeated 1000 times and compared to the actual diagonality metrics. Epoch segments that did not contain at least one output spike in all movement directions were dropped. An output unit was dropped if all three epochs were dropped, leaving 62/67 units for Monkey C and 58/78 units for Monkey N. All of the actual diagonality metrics were larger than the corresponding randomly generated diagonality metrics.

To test whether positive and negative contributions to the membrane potential along the diagonal balanced, the values of the diagonal were regressed against a cosine function. For both Monkey C and N, all remaining units had an input group/epoch combination with an *R*^2^*>*0.9. The cosine functions for all units were approximately centered on 0, showing that the positive and negative contributions were balanced.

## 5 Acknowledgments

We thank A. Whitford for collecting neurophysiological and behavioral data from Monkey C; D. Shi and X. T. Cui for electrode surface modifications of Utah arrays used with Monkey N. This work used Bridges-2 at Pittsburgh Supercomputing Center through allocation MED230028 from the Advanced Cyberinfrastructure Coordination Ecosystem: Services & Support (ACCESS) program, which is supported by National Science Foundation grants #2138259, #2138286, #2138307, #2137603, and #2138296. We acknowledge research grants from NIH (R01NS111148 to A.B.S.) and Pennsylvania HRFG (to A.B.S.).

## 6 Author contributions

A.B.S. conceptualized and supervised the project. H.M. and B.A.H. performed neurophysiological and behavioral data collection from Monkey N. H.M. processed and analyzed neurophysiological data. H.M. performed RNN training and analyses. A.B.S and B.A.H. performed SNN training and analyses. A.B.S. wrote the manuscript with input from all authors.

