## Supplementary Figures for "Hybrid Neural Network Models Explain Cortical Neuronal Activity During Volitional Movement"

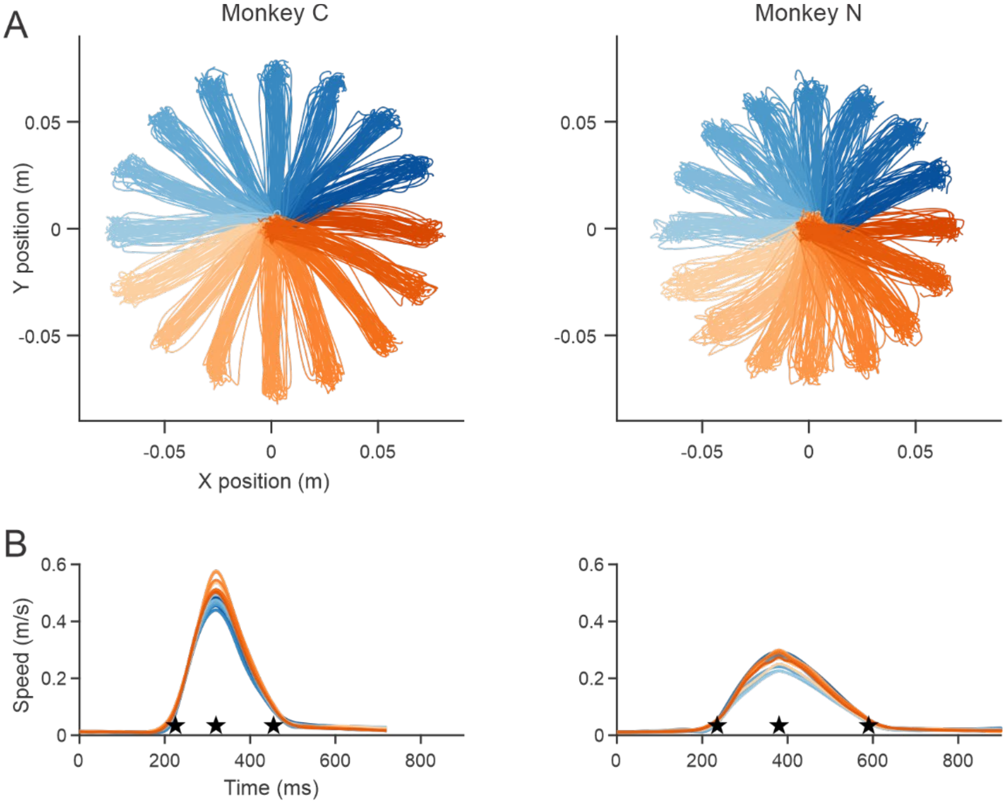


**Supplementary Fig. 1. Center-out reaching kinematics.** (**A**) Hand movement trajectories from all repetitions (47 for Monkey C and 49 for N) of reaching movements to 16 targets. Traces are color-coded by direction of the target. (**B**) Speed profiles averaged across trials for each target. The black stars indicate trial-averaged time of movement onset, peak velocity, and movement offset, respectively.

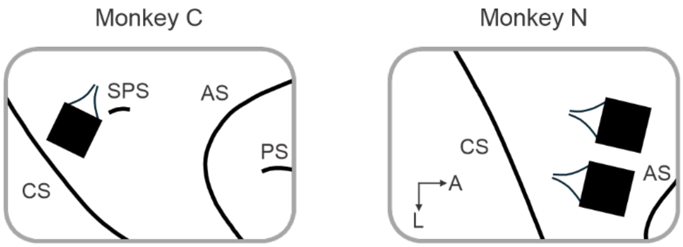


**Supplementary Fig. 2. Electrode array placement.** CS: central sulcus; SPS: superior pre-central sulcus; AS: arcuate sulcus; PS: principal sulcus. A: anterior; L: lateral. Black boxes represent Utah arrays (4x4 mm, drawn to scale). Small black lines connected to arrays represent wire bundles.

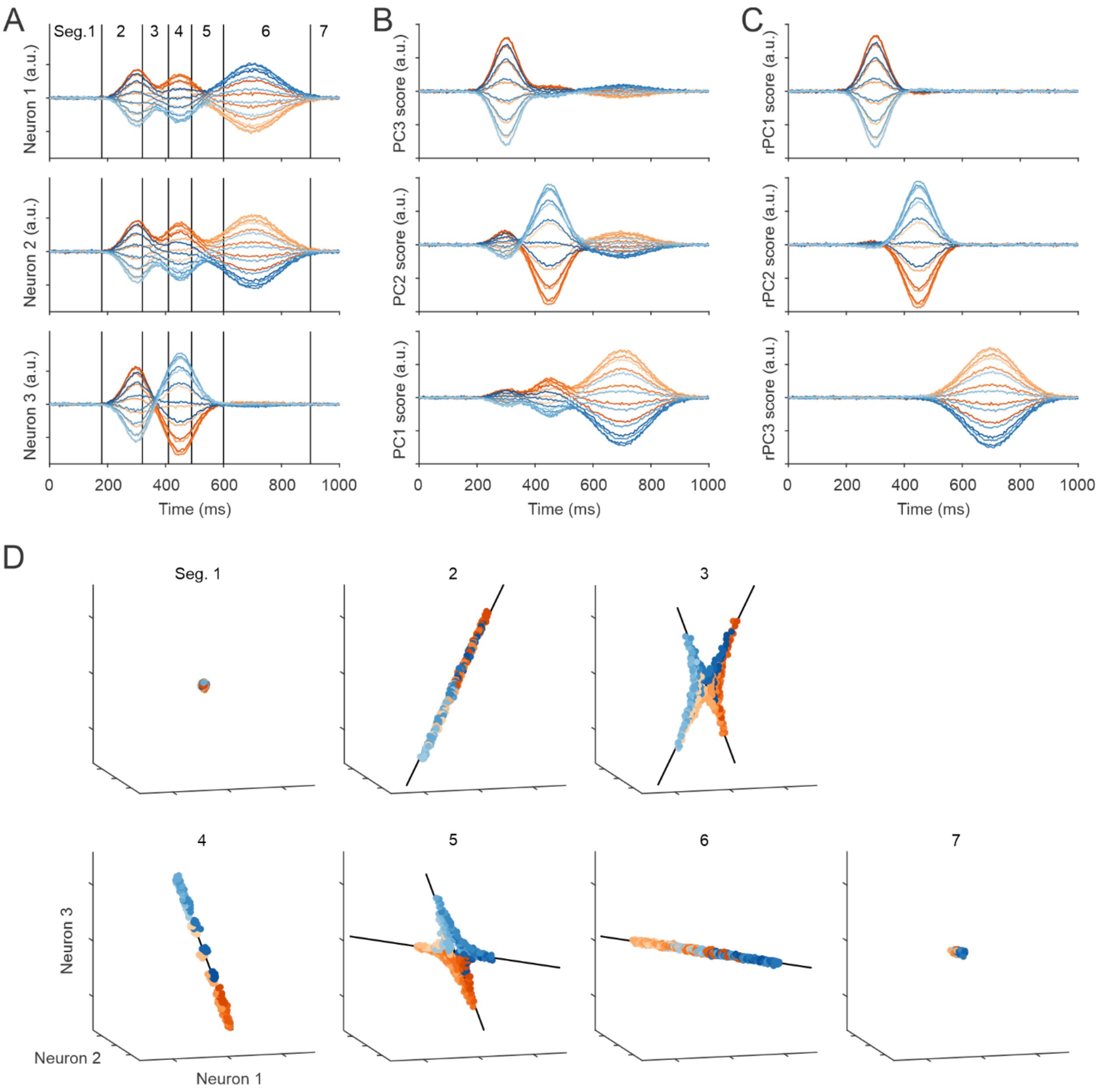


**Supplementary Fig. 3. An illustration of multiple states using three simulated neurons.** (**A**) Firing rates from three simulated neurons. Each colored line represents the trial-averaged firing rate during movements to one of 16 targets in a center-out task. Each neuron has 3 directionally tuned bell-shaped firing rate modulation epochs - states. The entire duration of simulation (1 sec) is divided into 7 segments, where there is mostly none (seg. 1 and 7), only one (seg. 2, 4 and 6), or two overlapping (seg. 3 and 5) firing rate modulation epochs. (**B**) PCA scores from the three simulated neurons. Scores are color-coded by movement direction. (**C**) Scores from PCA after Promax rotation (rPCA). Each of the three firing rate modulation epochs is uniquely captured by one of the three rPCA components. (**D**) Data clouds formed by sample points from each of the seven temporal segments defined in **A**. Black straight lines represent the rPCA axes.

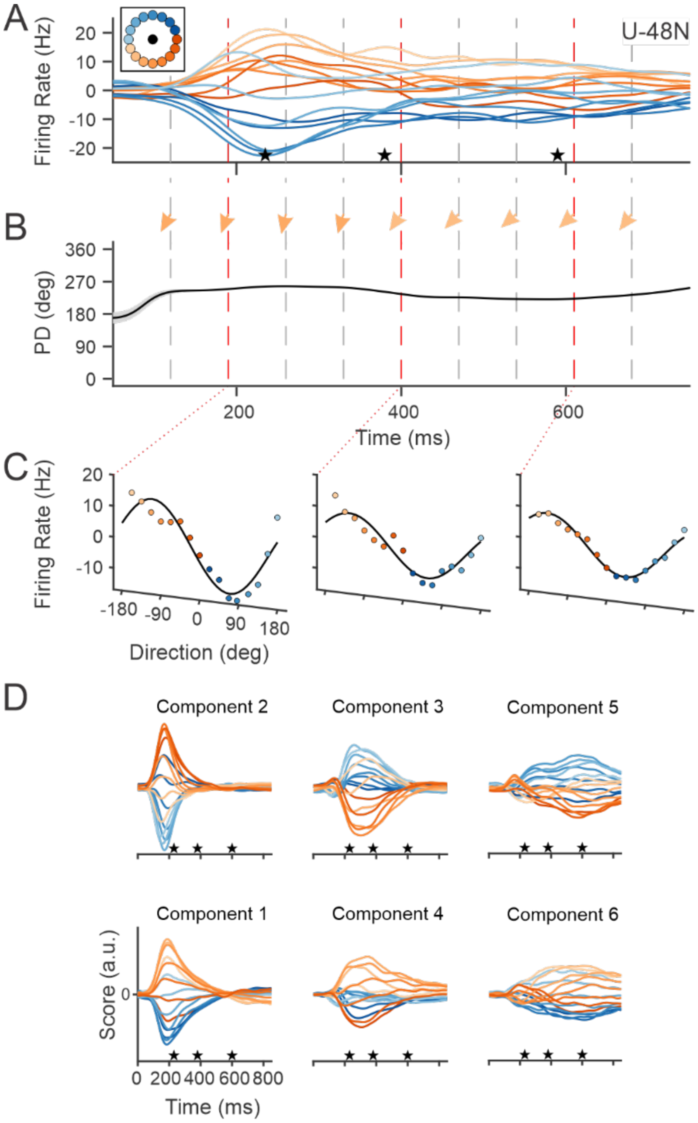


**Supplementary Fig. 4. Multiple states in a single reach.** (**A**) Histogram of firing rates from an example motor cortical unit (Unit 48, Monkey N). Each colored line is the trial-averaged firing rate during movements to one of 16 targets in a center-out task. The mean firing rate across all 16 targets at each time point was removed. (**B**) Preferred directions were calculated successively in 20 ms bins and are represented by the solid trace. Preferred directions are also symbolized by the arrows. (**C**) Three example tuning functions from points in the movement indicated by dashed lines and colored arrows in **A** and **B**. Firing rates are indicated by dots; solid lines are the cosine fits to the data. **(D)** rPCA scores from Monkey N. Scores are color-coded by movement direction. The profiles in each column have peaks at about the same time in the trial. Trials start from target onset, and are aligned to movement onset, peak velocity, and movement offset (indicated by the star markers). Reaction time is the period from time zero to the first star. Movements take place in the time span between the first and third star.

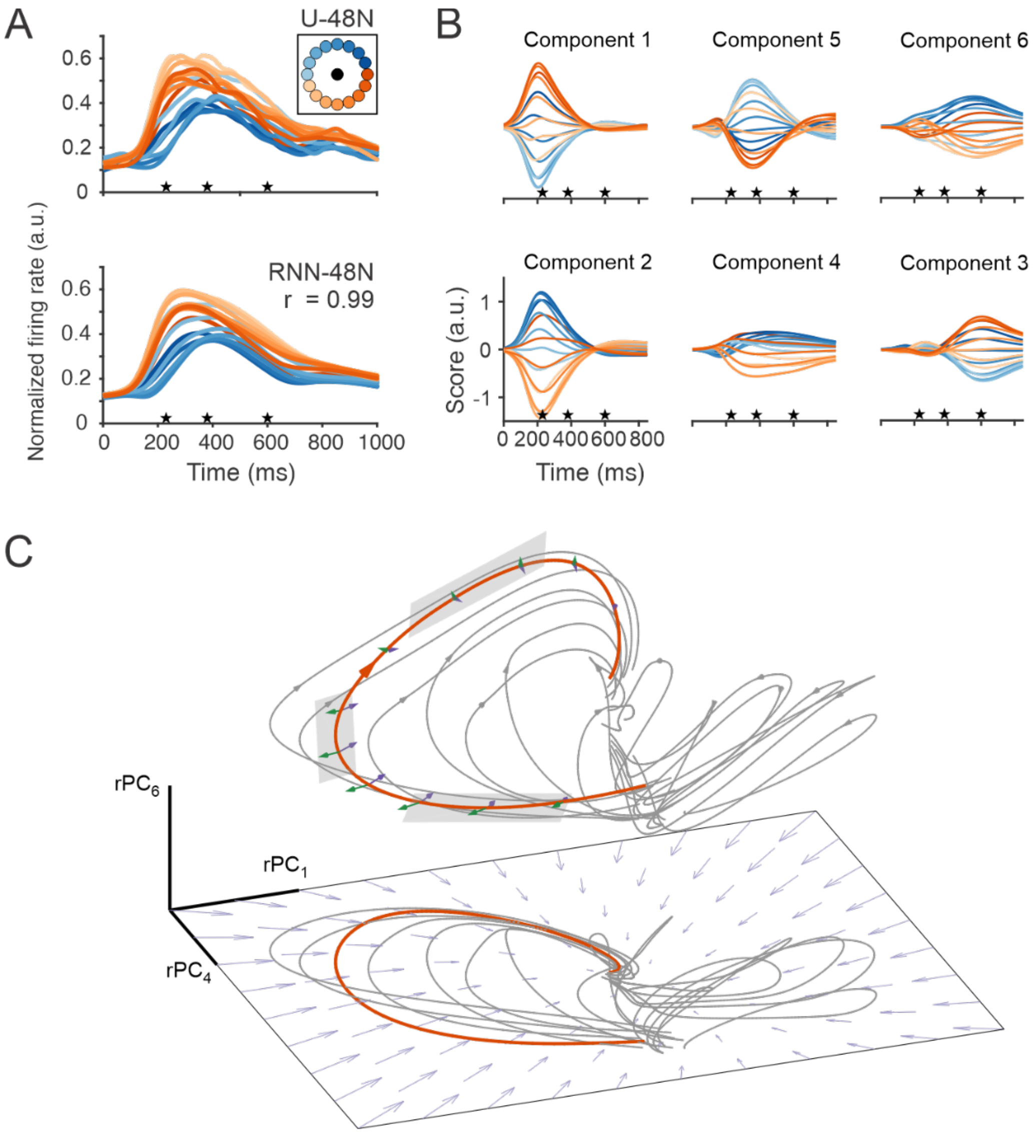


**Supplementary Fig. 5. Recurrent neural network – Monkey N.** (**A**) Actual (top) and predicted (bottom) firing rates of Unit 48 of Monkey N. Traces begin at target onset. Stars correspond to movement onset, peak speed, and the end of movement. The traces are color-coded by target direction. The actual-predicted firing rate correlation for this unit was 0.99. (**B**) rPCA scores of RNN units after training. The six highest scores (explained 94.6% data variance) are plotted after Promax rotation. (**C**) Neural dynamics of RNN in a 3-D subspace. Each axis corresponds to a rPCA component calculated from the activations of the RNN units. The 3-D space is formed with one dimension from each of the three pairs of rPCA components. Neural trajectories are plotted for sixteen trials by projecting RNN activations onto this 3-D subspace. Each trajectory starts from target onset and terminates at the end of target hold. Contributions to the trajectory gradient at evenly spaced locations are illustrated with purple (recurrent connections) and green (extrinsic input) arrows.

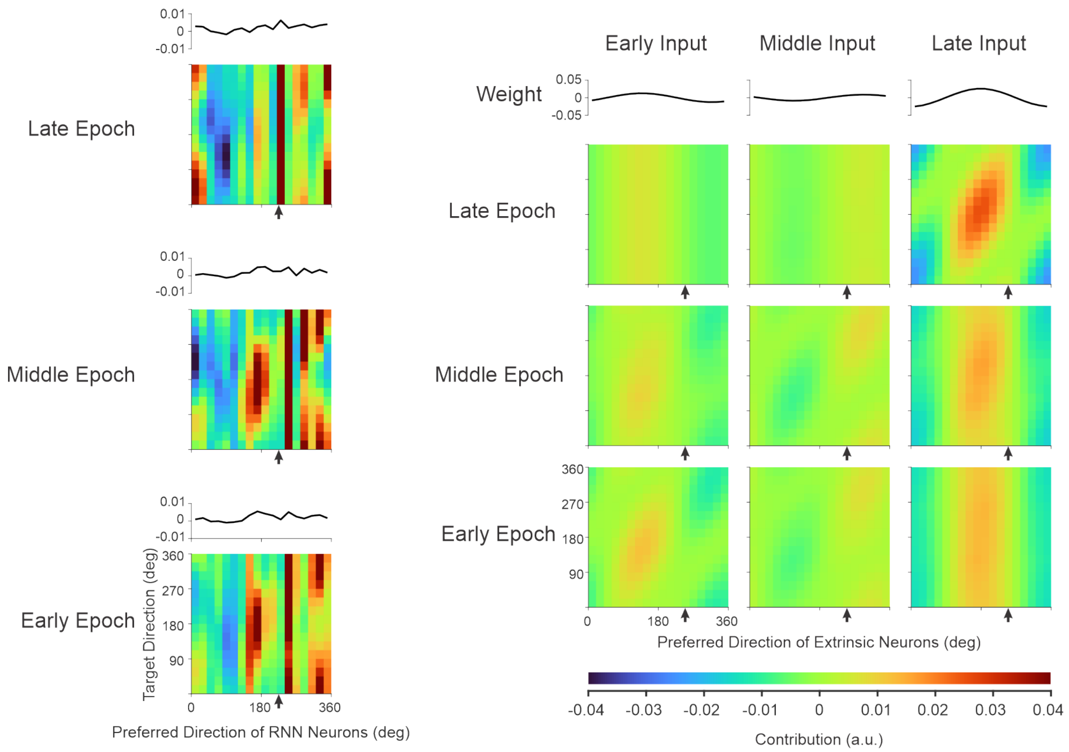


**Supplementary Fig. 6. Contributions to an example RNN unit – Monkey N example.** Contributions to the change in activation (x(t)) of RNN-48N. The contributions of the recurrent units (left) and the three input groups(right) are evaluated in the early, middle, and late epochs of the reaching movement. For each epoch, the RNN units are divided into 18 groups, according to their preferred direction (PD) in that epoch. The mean weights of the input group are plotted above each heat map. Each pixel of the heatmap shows the total contribution from the units in a PD group during an epoch of all trials with the same target location. The arrows below each weight profile designate the preferred direction of the output unit in the respective epoch.

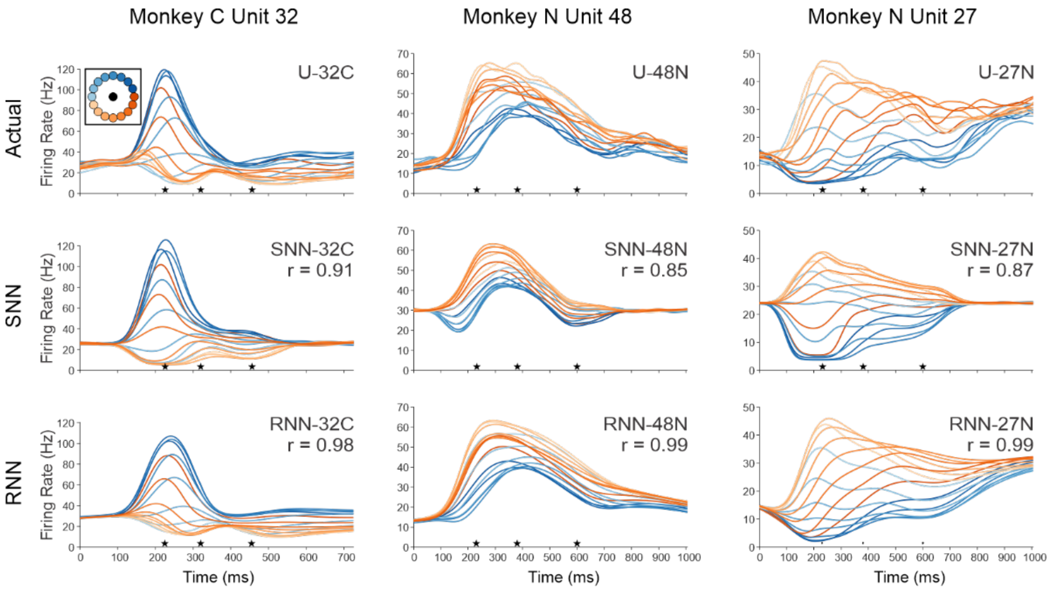


**Supplementary Fig. 7. Firing rate reconstruction by neural network models.** Actual (top) and predicted firing rates by SNN (middle) and RNN (bottom) of three example motor cortical units. Traces begin at target onset. Stars correspond to movement onset, peak speed, and the end of movement. The traces are color-coded by target direction. The r value is the correlation coefficient between actual and model generated firing rate.

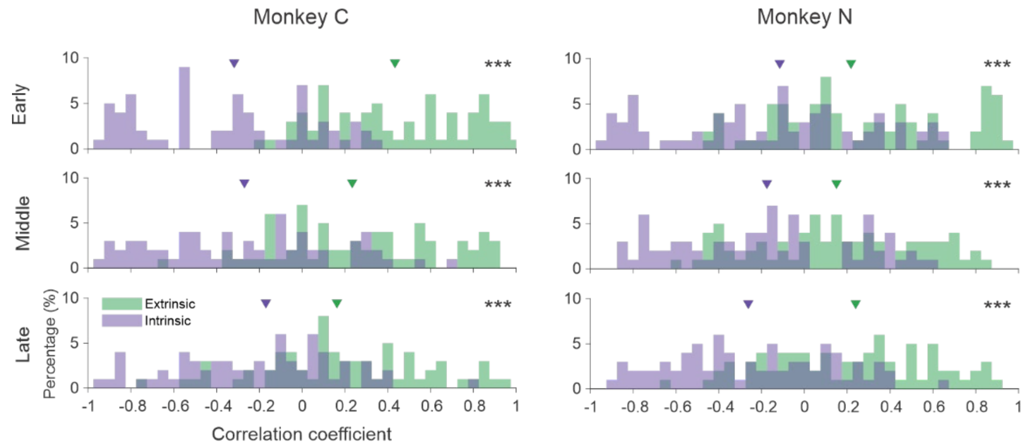


**Supplementary Fig. 8. Correlation between RNN output firing rate and its contributions.** Green: Distribution of correlation coefficients between extrinsic contribution and RNN unit firing rate across all output units. Purple: Distribution of correlation coefficient between intrinsic contribution and firing rate. The triangle markers represent the median values of the two distributions. The median correlation between extrinsic contribution and firing rate is significantly higher than that of the intrinsic contribution (one-tailed Wilcoxon signed-rank test, *** represents p<0.001) in all three temporal epochs.

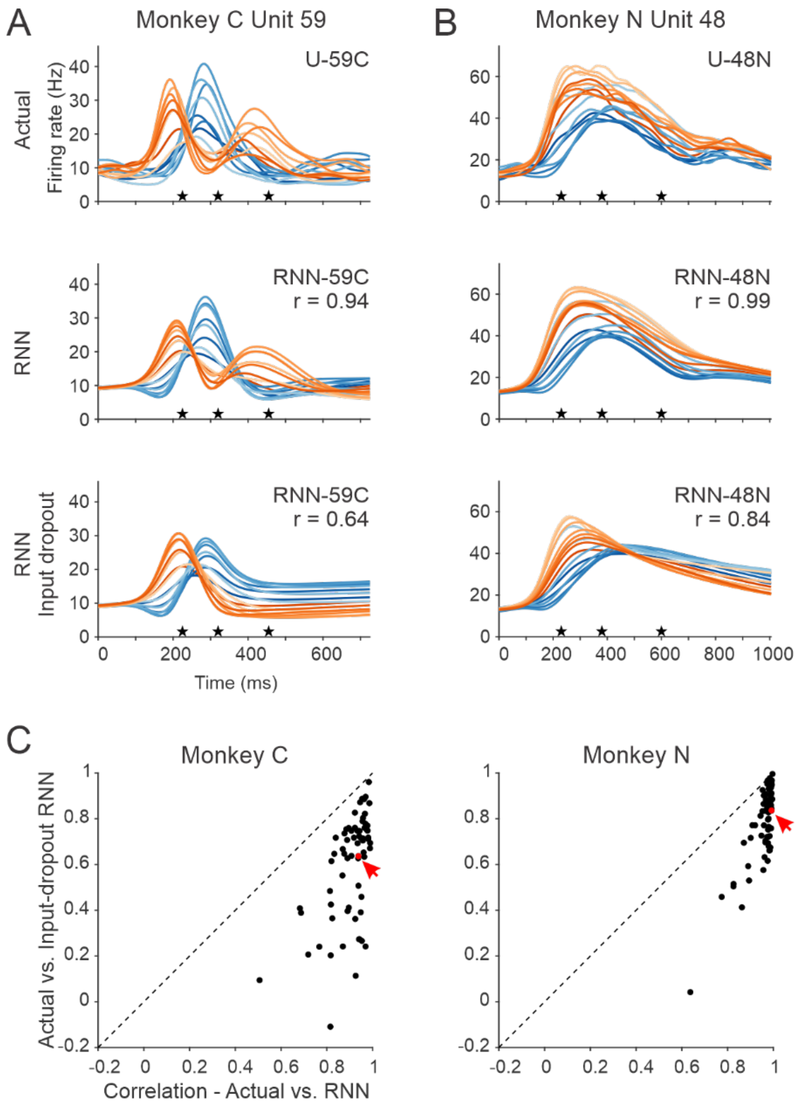


**Supplementary Fig. 9. Firing rate reconstruction by neural network models with input dropout.** (**A**) Top: Recorded firing rate profiles of Monkey C Unit 59. Middle: Predicted firing rates by RNN for this unit. The r value is the correlation coefficient between actual and RNN generated firing rate. Bottom: RNN predictions with the latter two groups of extrinsic inputs dropped out (see Methods). **(B)** Results for Monkey N Unit 48. **(C)** Correlation between actual and RNN predicted firing rate (abscissa) vs. correlation between actual and predicted firing rate by the RNN under input dropout (ordinate). Each dot represents one RNN output unit. The red dot is the example unit above.

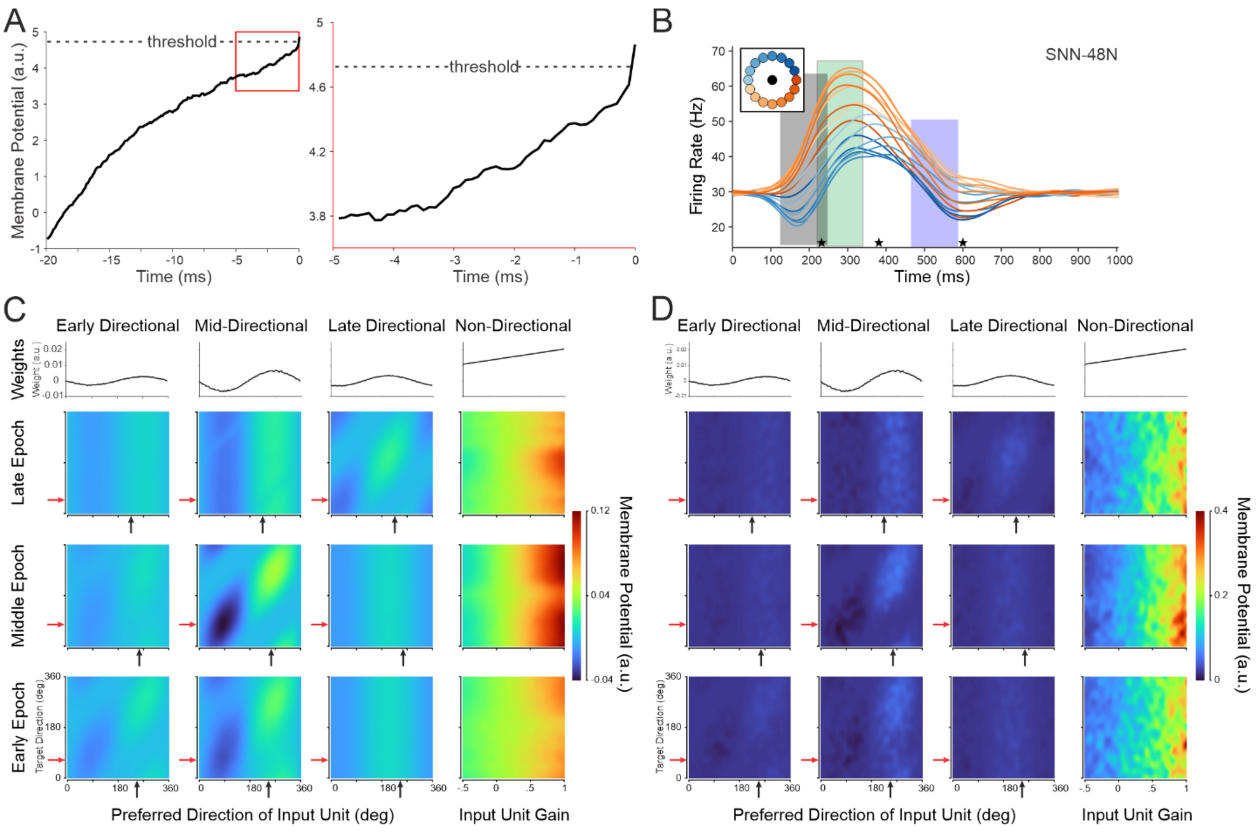


**Supplementary Fig. 10. Input contributions to output spiking – Monkey N example.** Contributions to SNN firing. **(A)** Spike-triggered average of membrane potential. The modeled membrane potential of Unit 48N before an action potential was averaged across all spikes of the Epoch 2 zone (green rectangle in **B**) for a reach at 112 degrees. (Left) 20 ms average before the spike. (Right) 5 ms before the spike. **(B)** Analysis zones for the three epochs defined by the rPCA analysis. The center of each zone was determined by the peaks of the PCA components and then shifted on each trial relative to the three behavioral landmarks (stars). **(C-D)** Heatmaps of the change in SNN-48N's membrane potential during the buildup **(C)** or for the trigger input **(D)**. Columns correspond to the input groups, the weights for each input group are shown in the top row, and the other rows correspond to the epochs shown in **B**. Black arrows show the epoch-specific preferred direction of SNN-48N. Red arrows show the anti-preferred direction.

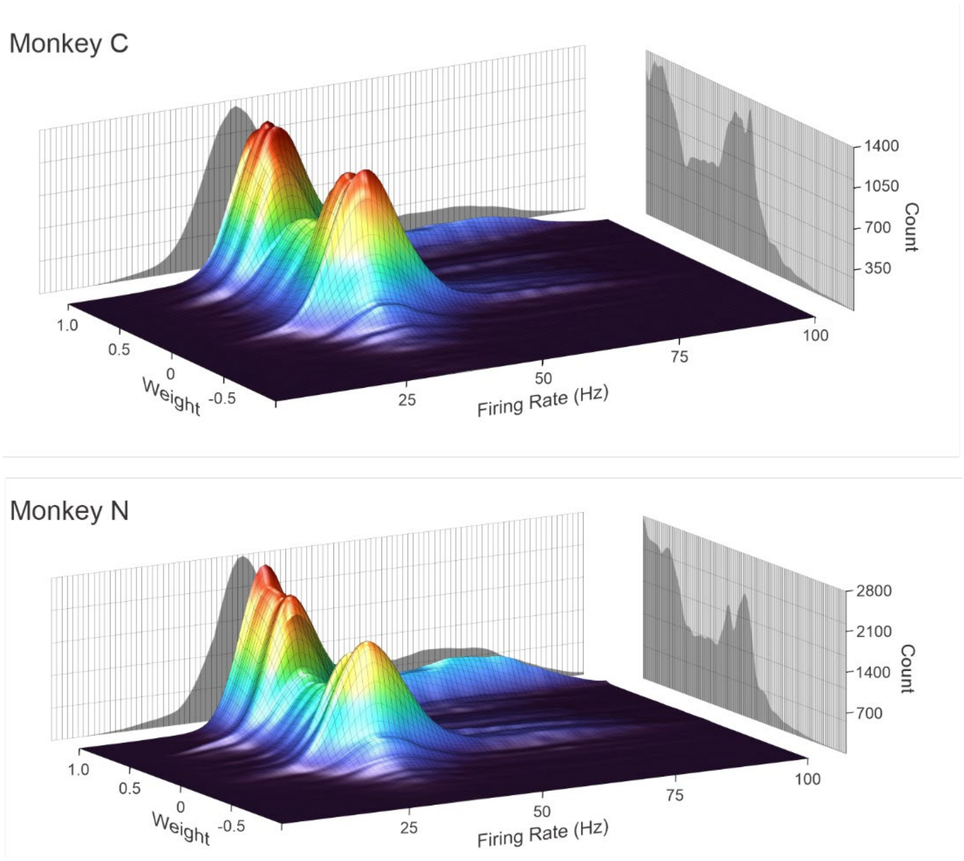


**Supplementary Fig. 11. Weights and firing rates of inputs *triggering* an output spike.** 2D histograms showing the total counts of input spikes from directional inputs in the trigger window before an output spike for all output neurons.

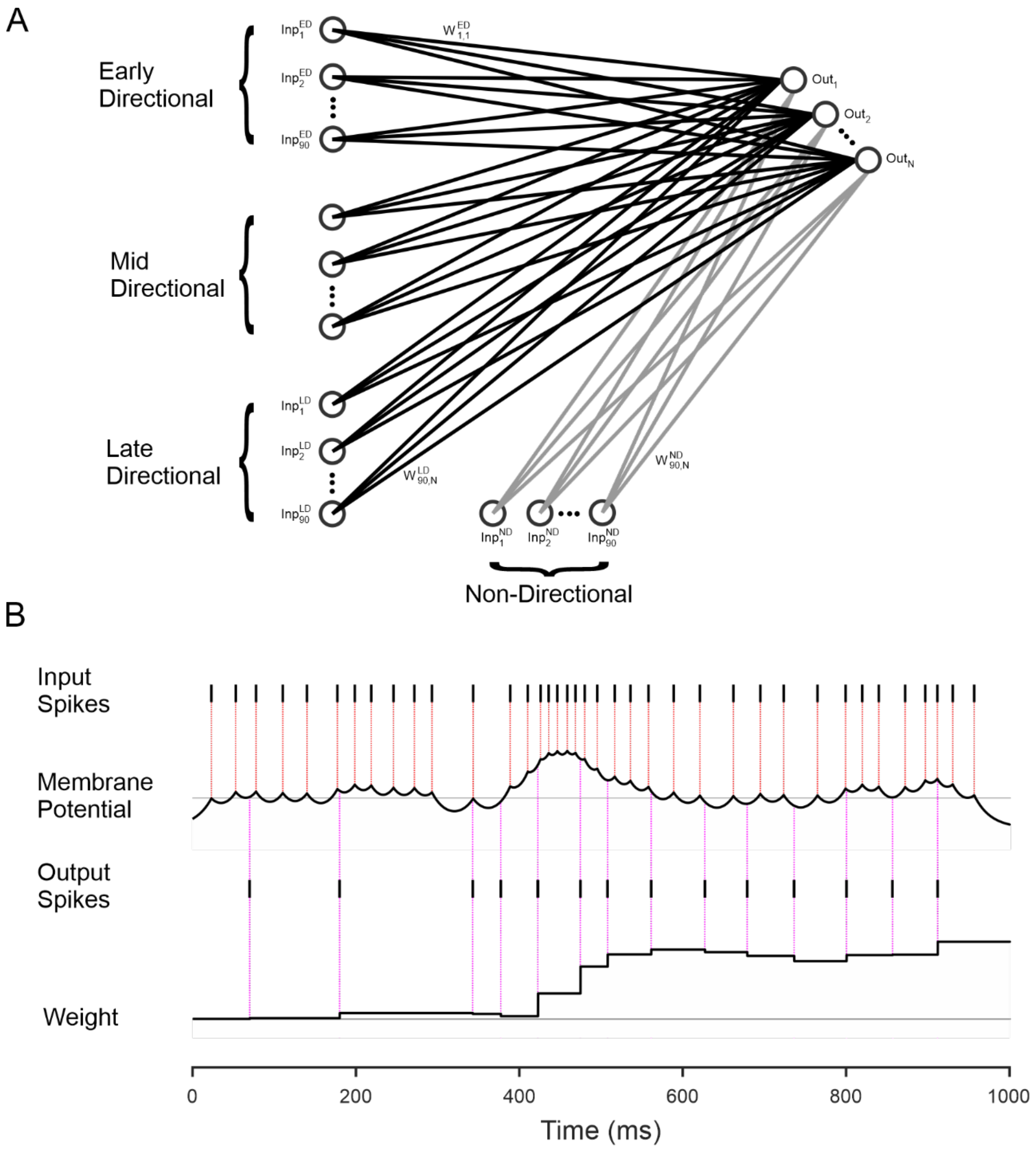


**Supplementary Fig 12. Spiking neural network.** **(A)** The four input groups, each with 90 neurons. Each input neuron (Inp) is fully connected to each output neuron (Out). Weights (W) are trained using a Hebbian learning rule. **(B)** Example trial showing how weights are trained – this process is repeated for each trial, and input/output neuron combination. Input spikes increase the membrane potential, which decays with a 20ms time constant. At the time of an output spike, the value of the membrane potential is added to the weight.
